# A unifying theoretical framework for tick-borne disease risk to explain conflicting results of exclosure experiments across scales

**DOI:** 10.1101/2024.10.03.616557

**Authors:** Ari S. Freedman, Simon A. Levin, Stephen A. Felt, Giulio A. De Leo

## Abstract

Disease ecology has focused greatly on determining how changes to biodiversity may drive infectious disease risk for humans. Fencing off experimental areas (exclosures) has been a common experimental approach to assess how removing large-bodied hosts may affect disease risk, especially with tick-borne pathogens (TBPs). However, exclosure experiments have found conflicting results based on the experiment’s scale, with smaller exclosures tending to increase tick densities inside the exclosure and larger exclosures tending to decrease tick densities inside. Previously, we have lacked a unifying theoretical framework able to reconcile the results of exclosure experiments across spatial scales. We present a spatially explicit model of TBP risk incorporating tick dispersal by small competent mammal hosts which can enter the exclosure and by large incompetent mammal hosts excluded from the exclosure. Our model reproduces the scale-dependence and spatial patterning observed in past exclosure experiments while elucidating their causal mechanisms. Specifically, the modeled exclosures produce high densities of infected ticks near their boundaries, with the densities decreasing towards the exclosure’s center. Empirical results have found lower tick densities at the exclosure’s edge than its center, a pattern we demonstrate can also be produced if we additionally allow ticks in their free-living questing stage to disperse.

## 1 Introduction

Few questions in the world of disease ecology have been as hotly debated as that of the relationship between biodiversity and infectious disease [1–18], and for good reason. In a time of rapid global declines in biodiversity and increases in infectious disease burden, it is crucial to understand how these changes have been interacting with each other in the hopes of using biodiversity maintenance as a means of disease control [18]. Furthermore, the diversity-disease relationship has been found to be highly contingent on a number of factors—including host species composition [3, 19–24], host species densities [25–29], pathogen life history [1, 2, 13, 18, 24], and the scale of measurement [16–18, 28, 30, 31] (the focus of this paper)—confounding any attempts at consensus.

Much of the biodiversity-disease debate has centered around tick-borne pathogens (TBPs), as tick and TBP life cycles are heavily affected by the assemblage of available host species, and TBPs such as the vectors of Lyme disease and tick-borne encephalitis have become increasingly common with anthropogenic climate change and human encroachment on tick-rich habitats [32–35]. At the simplest level, the diversity-disease relationship can be driven by either the “dilution effect” or the “amplification effect”. In the former case, increasing tick host species diversity dilutes tick feedings across a wider range of hosts some of which may be unable to transmit the pathogen, decreasing the disease risk to humans. In the latter case, increasing tick host species diversity can amplify tick abundance and ultimately increase the disease risk to humans [36, 37]. Experiments using exclosures—fenced-off areas built to keep out larger animals, which are often the most affected by declining biodiversity [38]—have been the primary tool of experimental manipulation to assess how the extirpation of large-body size hosts may affect TBP risk for humans.

However, exclosure experiments have notoriously produced conflicting results: some studies, typically associated with small exclosures [39–41], have found that removing large incompetent hosts increased the densities of infected nymphs (the life stage that poses the greatest TBP risk to humans), indicative of dilution. Other studies, typically associated with larger exclosures [42, 43], have found the opposite, that removing large incompetent hosts decreased densities of infected nymphs, suggesting amplification [31]. Scale-dependence has been proposed as a potential explanation for the conflicting results from exclosure experiments and studies of diversity-disease relationships in general [16–18, 31]. Previous work has hypothesized that this scale-dependence can occur when the large mammals being excluded are necessary for the tick’s life cycle but are also incompetent TBP vectors [31, 44], as is the case with deer in many TBP systems [22, 31, 45] (we will use deer and rodents as the motivating examples of large and small mammals, respectively, throughout this paper). In this scenario, small exclosures can produce higher infected nymph densities due to the lack of deer, which would otherwise take up ticks from the environment while blocking the TBP transmission cycle. In large exclosures, by contrast, the central area of the exclosure may become completely void of ticks due to the lack of deer in the surrounding area and the limits of rodent-based tick dispersal, leading to a lower density of infected nymphs in the exclosure on average

While previous work has discussed conceptually how exclosure size may affect the outcome of these manipulative experiments [6, 9, 16, 17, 31], there is a paucity of theoretical models that reconcile the apparently conflicting results of exclosure experiments over multiple scales. To our knowledge, there is only one theoretical study that has attempted this, approximating space with a simplified two-patch model representing the inside and the outside of the exclosures as homogeneous areas [44]. However, this approach is unable to explicitly account for exclosure size, and ignores spatial gradients in disease risk that may be present within an exclosure.

Here we present a unifying, spatially explicit theoretical framework incorporating tick and pathogen life cycles and dispersal via rodent and deer hosts to explain the scale-dependence observed in exclosure experiments. While previous exclosure experiments and models of theoretical exclosure experiments only measure average tick densities inside and outside the exclosure, our approach maps tick densities at every point within the two-dimensional area being modeled. This added nuance reveals stark decreasing gradients in infected nymph density going from the edge of an exclosure to its center, highlighting that TBP systems may be under the effects of both dilution and amplification at different points in space. Specifically, our results show that tick-borne disease risk is highest at the exclosure’s edge where deer are nearby, while disease risk is lowest at the exclosure’s center far away from any deer, as deer are required to support the tick population. Thus, we emphasize the need for exclosure experiments to empirically test for gradients in tick densities within exclosures to shed light on the spatial dependence of the mechanisms at play and the role of host movement.

Our spatially explicit modeling approach also allows us to incorporate movement by ticks in their free-living “questing” stages, which we show can produce gradients that are either non-monotonic or reversed from what would be expected without questing tick movement. These results with questing tick movement match up well with the spatial patterns found by the few studies that have reported on tick gradients within exclosures [14, 16, 31]. We also derive mathematical approximations to the gradients that result from our model, proving that non-monotonic gradients in infected nymph density can only result when there is questing tick movement and a sufficiently high rate of contact between nymphs and deer. This result lends credibility to the “tick sharing” hypothesis [16, 31] stating that tick densities near the inside edge of the exclosure should be lowered by questing tick movement as ticks are able to move outside the exclosure and picked up by deer.

## 2 Model overview

We modeled a system with one species of three-host tick, one TBP, and two tick host types, rodents and deer. Thus, each tick feeds on three hosts throughout its life, each of which is either a rodent or a deer, and each blood meal corresponds to a different tick life stage (larva, nymph, and then adult). Though all possible hosts for the tick are lumped into one of two categories (“rodents” representing all small mammal hosts and “deer” representing all large mammal hosts), ticks in reality may feed on a wide variety of mammal and non-mammal hosts. Drawing on other TBP-modeling literature [46–50], we utilized a mechanistic movement modeling approach that models tick and host densities at each spot in a two-dimensional area over time using spatial partial differential equations to represent host movement (and questing tick movement in Section 3.2) as undirected Brownian motions. Equivalently, the densities at each spot in space and time follow a system of ordinary differential equations (Fig. 1*b*), while each host (and questing tick in Section 3.2) also independently moves randomly through space (Fig. 1*a*).

**Figure 1:**
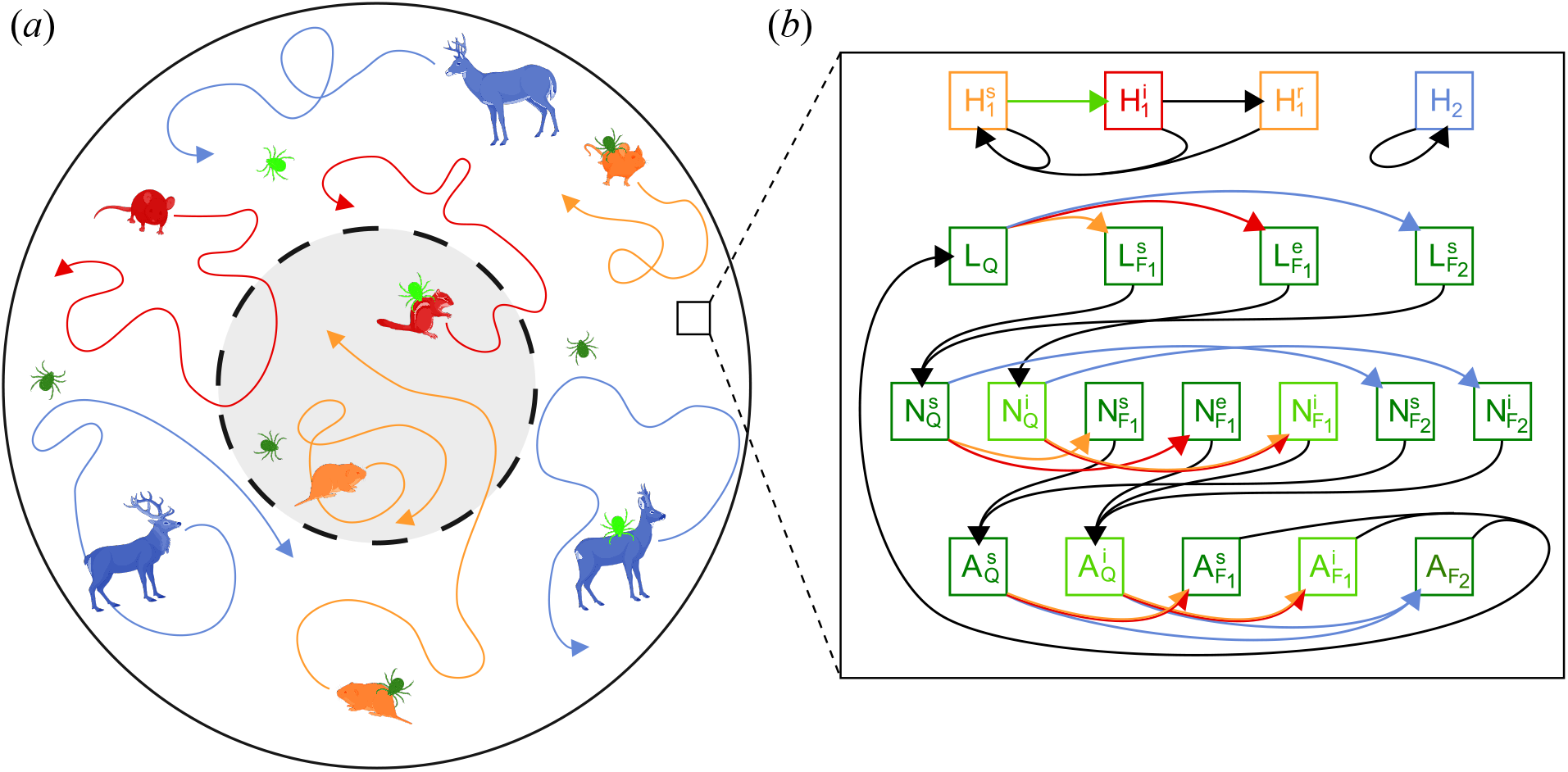
Diagram of our mechanistic movement model, consisting of (*a*) the movement component and (*b*) the ordinary differential equation component. In (*a*), deer and rodent hosts are constantly moving in Brownian motions with deer restricted from the exclosure area (gray circle with dash boundary), while ticks are either feeding on a host and moving with them or questing and stationary (in Section 3.2 we also incorporate questing tick movement). At each point in space and time, tick and host densities are controlled by a system of ordinary differential equations, shown in (*b*). The four rows of compartments represent hosts (*H*_1_ for rodents, *H*_2_ for deer), tick larvae (*L*), tick nymphs (*N* ), and tick adults (*A*), respectively. Superscripts represent infection status (*s, e, i*, or *r* for susceptible, exposed but not infectious, infectious, and recovered, respectively), while subscripts for ticks denote their feeding status (*F*_1_ for feeding on rodents, *F*_2_ for feeding on deer, and *Q* for questing). Colored arrows indicate a transition instigated by interaction with a different animal category: blue for deer, red for infectious rodents, orange for non-infectious rodents, light green for infectious ticks, and dark green for non-infectious ticks. See supplement for full model description. Diagram made with BioRender.

The ordinary differential equation component of the model (Fig. 1*b*) and its associated parameters were adapted from previous work by Pugliese and Ros`a [25, 44] modeling tick-borne encephalitis in a tick-deer-rodent system, though it can be easily parameterized to represent other TBP systems. The tick population is structured in three life stages (larva, nymph, or adult) which vary in their infection status (susceptible, exposed, or infectious) and whether they are actively feeding (on a rodent or deer) or questing. Larvae and nymphs able to feed on either rodent or deer but adults feeding almost exclusively on deer [25, 31]. We modeled rodents as competent TBP vectors, while deer are incompetent TBP vectors that can contribute to the tick’s life cycle but are unable to transmit the pathogen [22, 31, 45]. Accordingly, rodent hosts exhibit susceptible-infected-recovered dynamics with lifelong immunity and infection triggered by being fed upon by an infected tick, while all deer are lumped into one category. Ticks are all born as susceptible larvae and can become infected in subsequent life stages, either by feeding on infected rodent hosts or by sharing a host with other infected ticks (co-feeding transmission [51, 52]), after which they stay infectious for the rest of their lives.

For the spatial component (Fig. 1*a*), we considered a circular domain with a radius of 1 km over which the model is simulated, with a smaller concentric area inside representing the exclosure. Though most exclosures are rectangular in practice, the circular shape ensures radial symmetry, effectively reducing the dimensionality to one to make simulations more efficient. Tick and host dynamics and movement are identical inside the exclosure, except that no deer are present inside the exclosure. Thus, rodents randomly roam in Brownian motions unrestricted throughout the larger circular domain, while deer also move around randomly but only outside the exclosure. In Section 3.2, we additionally assumed that free-living questing ticks can move in Brownian motions in between feedings on hosts. Rodents suffer infection-based mortality, so their population size may vary across space and time. For deer, they do not suffer infection-based mortality so their population size becomes constant across space and time; however, it is still important to track deer movement for the sake of the ticks feeding on them.

We ran simulations for 3,000 days to get maps of equilibrium infected tick densities over space, considering exclosures with radii up to 300 m. All simulations start with tick and host densities constant over space at a pre-exclosure equilibrium. We parameterize rodent diffusion so that a single rodent moves on average 10–50 m/day, while questing ticks are able to move an average of 5 m/day in Section 3.2. Full model equations and parameters can be found in the supplement.

## 3 Results

### 3.1 Model without questing tick movement

By modeling space explicitly, we are able to parameterize exclosure size directly and predict how disease risk within the exclosure will vary as a function of exclosure size. In Fig. 2, we show the average equilibrium densities of infected questing nymphs inside and outside the exclosure for a range of exclosure sizes and for three different rodent movement rates. The average infected questing nymph density is significantly higher inside the exclosure than it is outside when the exclosure is small, while for large exclosures the opposite is true. As exclosure size increases, the average infected questing nymph density inside the exclosure steadily declines, while the average density outside stays relatively flat.

**Figure 2:**
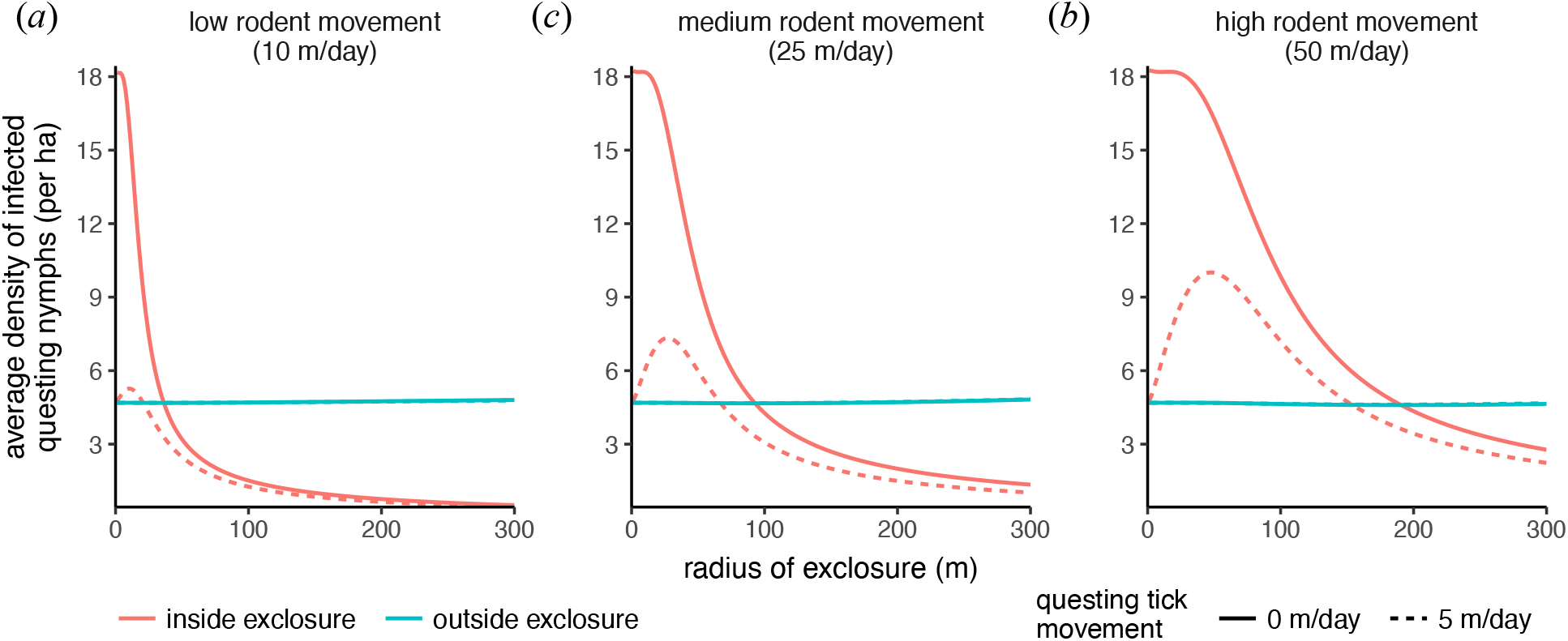
Density of infected questing nymphs at equilibrium, averaged across the inside of the exclosure (pink) and the outside of the exclosure (blue), for circular exclosures of varying size up to 300 m in radius. Solid lines show the equilibrium densities when there is no questing tick movement, and dashed lines when questing ticks move an average of 5 m/day (the blue dashed lines largely coincide with the blue solid lines). Rodents move an average of 10 m/day in (*a*), 25 m/day in (*b*), and 50 m/day in (*c*).

Our modeling approach also allows us to vary the rodent movement rate independently of exclosure size. The exclosure size at which the average infected questing nymph density is equal inside and outside the exclosure (we call this the “inflection exclosure size”) increases with rodent movement rate (Fig. 2): the inflection exclosure size is 0.4 ha (36.3 m radius) when rodent movement is low (10 m/day, Fig. 2*a*), 2.7 ha (92.8 m radius) when rodent movement is intermediate (25 m/day, Fig. 2*b*), and 11.4 ha (190.6 m radius) when rodent movement is high (50 m/day, Fig. 2*c*). This pattern is summarized in Fig. 3, where we can also see that the inflection exclosure radius is approximately linearly proportional to rodent movement rate, with the slope of this relationship increasing with higher tick-host-contact rates.

**Figure 3:**
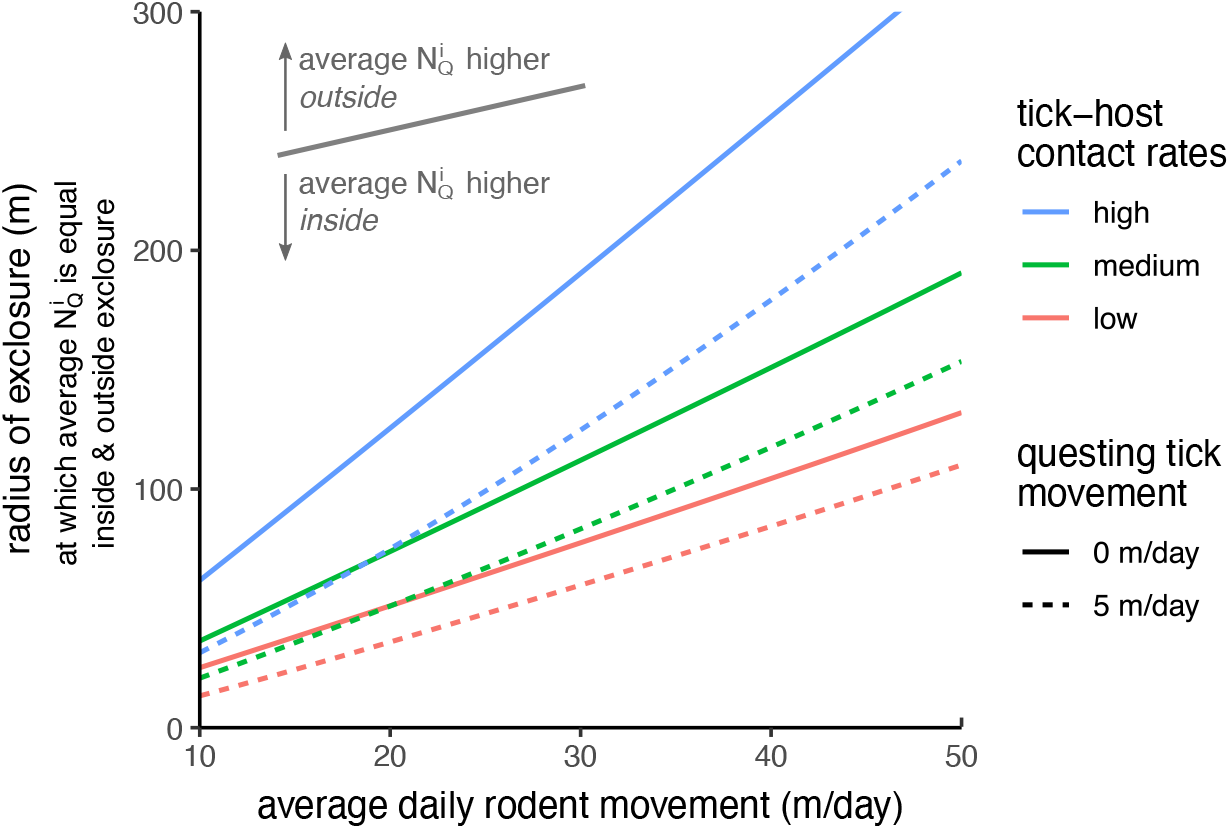
Exclosure radius at which the average density of infected questing nymphs 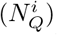 is equal inside and outside the exclosure, as a function of rodent movement rate. Each curve represents a different parameter set with either high (blue), medium (green) or low (green) tick-host contact rates, and questing ticks either having no movement (solid) or moving an average of 5 m/day (dashed). For a given parameter set and its corresponding curve, a point above the curve represents an exclosure size and rodent movement rate for which the average infected questing nymph density is higher outside the exclosure than inside, and the opposite is true for points below the curve. All other figures in this paper use the “medium” tick-host contact rates shown here.

Our spatially explicit model can also map TBP risk to humans at every point in two-dimensional space (Fig. 4), both at equilibrium (Fig. 4*a*–*b*) and throughout time (Fig. 4*c*–*d*). Doing so provides a clearer picture of the mechanisms that produce the scale dependence our model and past exclosure experiments have found. Specifically, we see a declining gradient in infected questing nymph density as we move from the edge of the exclosure to the center, for both small (50 m radius, 0.8 ha, Fig. 4*a*) and large (150 m radius, 7.1 ha, Fig. 4*b*) exclosure sizes, due to the limits of tick dispersal. Because questing ticks cannot disperse on their own, they rely on rodents to carry them from the outside, where almost all tick larvae are born due to the adults’ dependence on deer. As a result, tick densities in the exclosure are highest at the edge and declining towards the center. At the same time, the lack of deer, which could otherwise remove ticks from the environment and block TBP transmission as incompetent hosts, uniformly increases the questing nymph densities inside the exclosure, especially for infected questing nymphs. This leads to a strong dilution effect at the edge of the exclosure as the lack of deer increases infected questing nymph densities, while in larger exclosures the center is dominated by an amplification effect as the lack of deer and tick dispersal causes ticks to be nearly absent (Fig. 4*b*).

**Figure 4:**
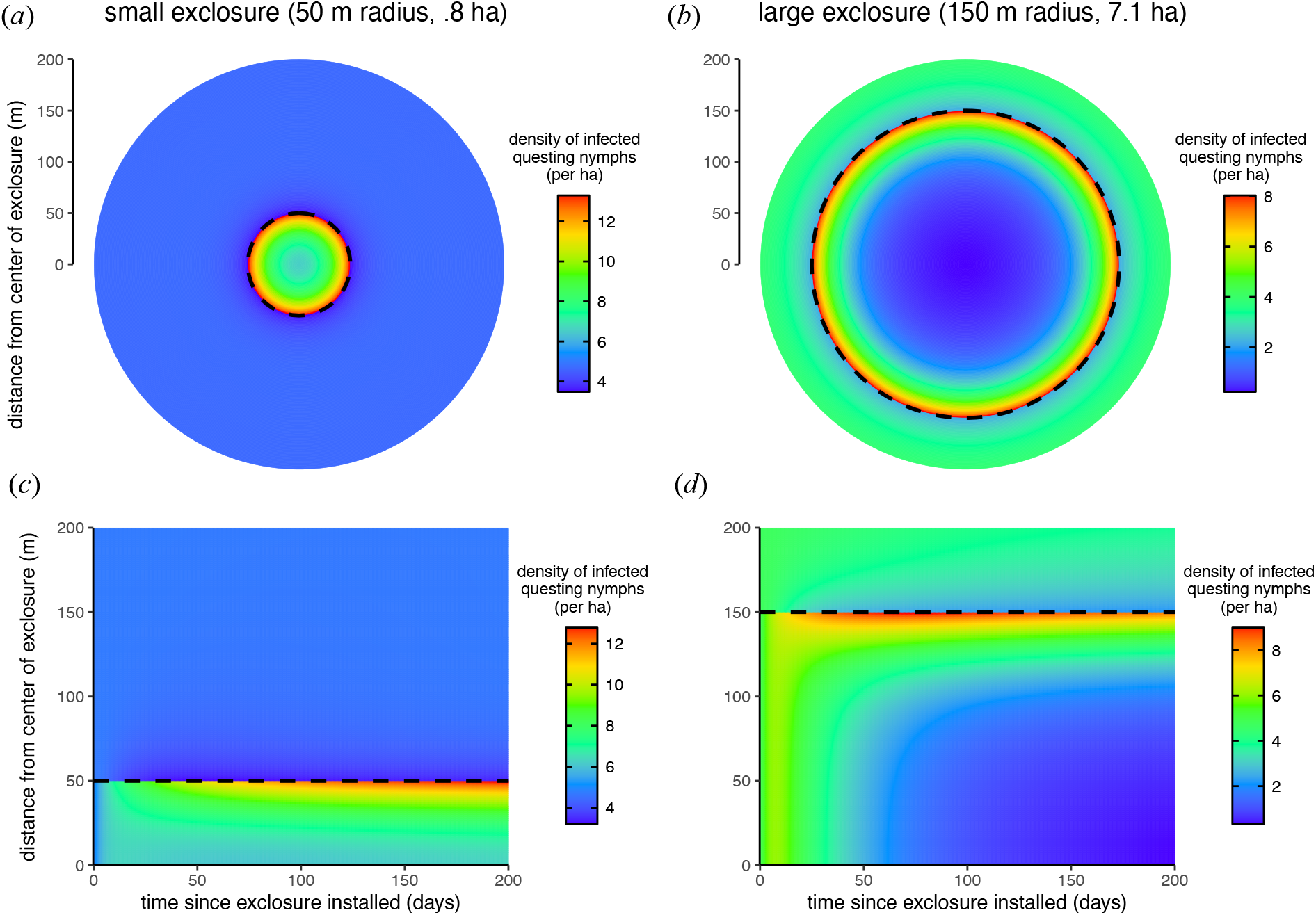
Infected questing nymph density across space, at equilibrium in (*a*)–(*b*) and over time in (*c*)–(*d* ). The exclosure is represented by the dashed black circle/line, with (*a*) and (*c*) modeling a small exclosure (50 m radius, .8 ha), (*b*) and (*d* ) modeling a large exclosure (150 m radius, 7.1 ha). In (*c*)–(*d* ), distance from the center of the exclosure is represented along the *y*-axis, since the model is radially symmetric, and the model starts from a pre-exclosure equilibrium which is constant across space. Note that the color scale bars in (*a*)–(*d* ) are all different, in order to emphasize the gradients in infected questing nymph density that exist within the exclosures. These simulations use intermediate rodent movement (25 m/day) and no questing tick movement.

If we make the simplifying assumptions that rodents suffer no pathogen-based mortality and adult ticks do not feed on rodents (both of which are true or close to being true in many systems [25, 31, 53]), and choose the exclosure to be the positive *x*-coordinate half of the plane (approximating a very large circular exclosure), we can analytically derive the gradient of infected questing nymph density 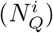 inside the exclosure at equilibrium:

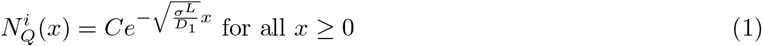

where *x* is the distance from the exclosure’s edge, *σ*^*L*^ is the rate at which feeding tick larvae fall off of their hosts (i.e. 1*/σ*^*L*^ is the average amount of time a larva spends feeding), *D*_1_ is the diffusion coefficient for rodent movement (equal to 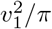 where *v*_1_ is a rodent’s average daily movement), and *C* is a positive constant. In other words, the infected questing nymph gradient follows an exponential decline at rate 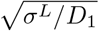 as we move into the exclosure. See the supplement for the full derivation.

### 3.2 Model with questing tick movement

Now we investigate the model’s behavior when we allow questing ticks to move independently of their deer or rodent hosts, in addition to their host-facilitated movement. For this section, we have questing tick nymphs and adults moving randomly in Brownian motions with an average daily movement of 5 m, while questing larvae still do not move due to their relatively limited mobility.

Like the model without questing tick movement, we still see that average equilibrium density of infected questing nymphs is equal inside and outside the exclosure for just one non-zero exclosure size, and this inflection exclosure size still increases approximately linearly with rodent movement rate (Figs. 2–3). However, questing tick movement uniformly lowers the average equilibrium density of infected questing nymphs inside the exclosure for all exclosure sizes, while the average density outside stays relatively unchanged (Fig. 2). Consequently, the inflection exclosure size is also lowered by adding questing tick movement (Fig. 3). This reduction in infected questing nymphs densities is most extreme for small exclosures, for which the average densities inside and outside the exclosure approach equality as the exclosure is shrunk, creating a non-monotonic relation between exclosure size and average infected questing nymph density inside the exclosure (Fig. 2).

The reason why questing tick movement reduces questing tick densities inside the exclosure actually lies outside the exclosure: while the only way for stationary questing ticks inside the exclosure to leave their questing stage is to either die or be picked up by rodents, mobile questing ticks may additionally move outside the exclosure and be picked up by deer, thereby increasing the rate at which they leave the questing stage, especially for ticks near the exclosure’s edge. Thus, we should expect the contact rate between questing nymphs and deer to have a significant impact on the way questing tick movement affects infected questing nymph densities across space.

In fact, with the high nymph-deer contact rate we use in these simulations, we see that the reduction in infected questing nymph densities near the exclosure’s edge is so strong it produces gradients in the exclosure that are either non-monotonic (in large exclosures, Fig. 5*b*) or completely reversed (in small exclosures, Fig. 5*a*) from how they are without questing tick movement (Fig. 4*a*–*b*). In the supplement, we show analytically that this phenomenon of non-monotonic or reversed gradients in infected questing nymph density can only appear when there is questing tick movement and the nymph-deer contact rate is sufficiently high. We support this with simulations showing that when the nymph-deer contact rate is low, the gradients inside the exclosure retain their usual monotonically decreasing nature even with questing tick movement—although this may come at the loss of the dilution effect as deer no longer provide their service of diluting nymph infection risk (Fig. S1).

**Figure 5:**
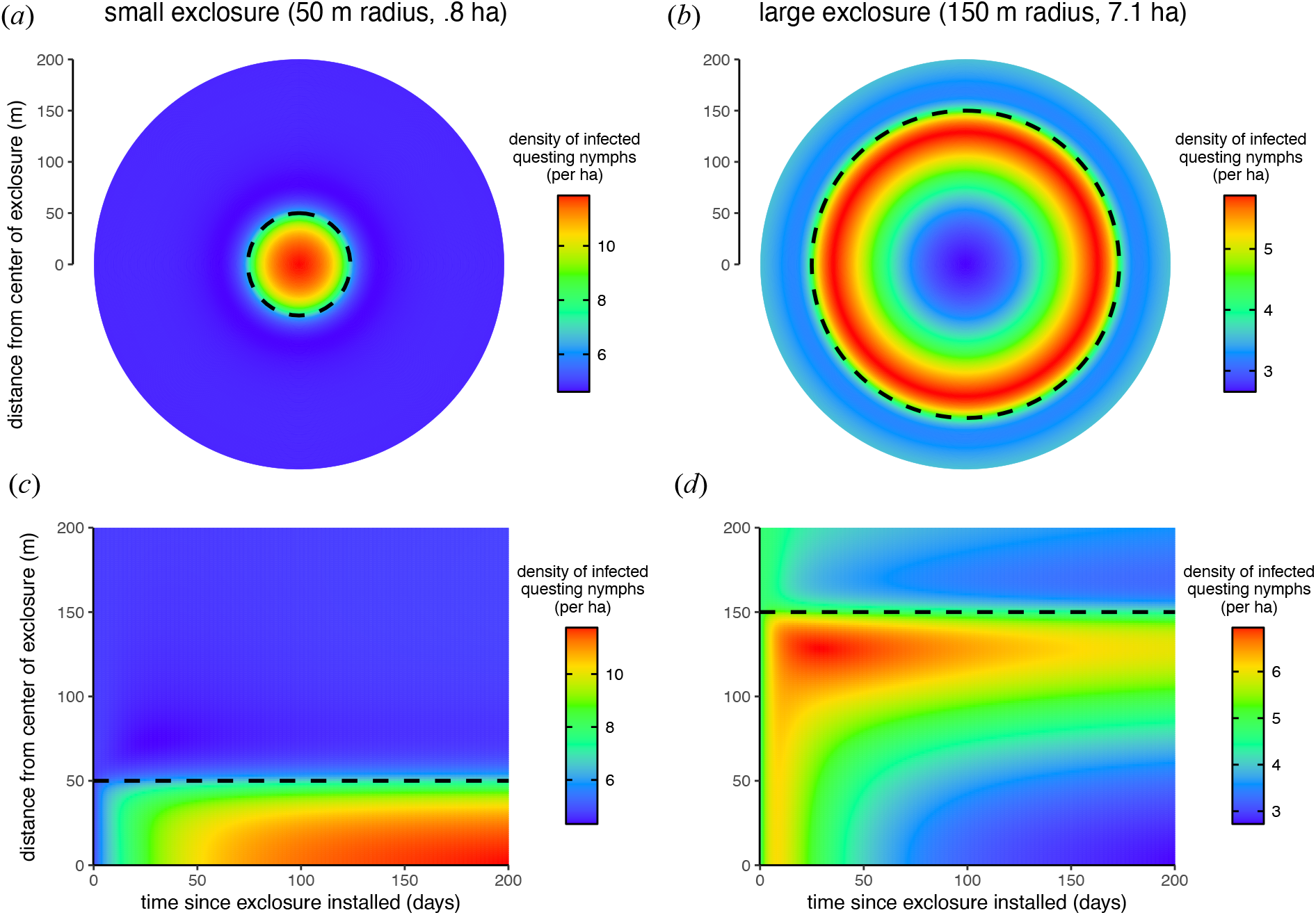
Everything is the same as for Fig. 4, except these simulations are run with questing tick movement (average of 5 m/day) and high rodent movement (average of 50 m/day). Questing tick movement can create infected questing nymph density gradients inside the exclosure that are either non-monotonic or reversed from the gradients produced without questing tick movement. Note that the color scale bars in (*a*)–(*d* ) are all different, in order to emphasize the gradients in infected questing nymph density that exist within the exclosures.

We also show in the supplement that for large exclosures, when there is no pathogen-based mortality or adult ticks feeding on rodents, the gradient’s rate of exponential decline as we move away from the exclosure’s edge approaches min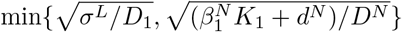 , or

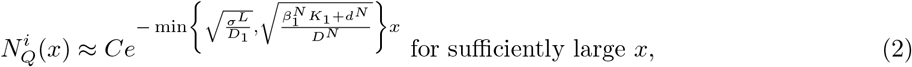

Where 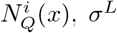, *D*_1_, and *C* are the same as for (1), *D*^*N*^ is the diffusion coefficient for questing nymph movement, 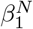 is the contact rate between nymphs and rodents, *K*_1_ is rodents’ carrying capacity density, and *d*^*N*^ is the death rate of questing nymphs. Since rodents have much higher movement rates than questing ticks (*D*_1_ ≫ *D*^*N*^ ), it should generally be true that 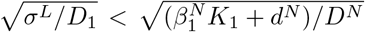; thus, questing tick movement likely does not change the fact from (1) that the gradient’s asymptotic rate of exponential decline is 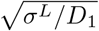. Either way, the nymph-deer interaction rate has no bearing on the gradient’s exponential decline away from the exclosure’s edge, despite its importance for determining the gradient’s pattern near the exclosure’s edge.

The transition in infected questing nymph density from the inside edge of the exclosure to the outside edge is much smoother and less dramatic with questing tick movement (Fig. 5), as opposed the sharp discontinuity in densities we see at the exclosure’s edge without questing tick movement (Fig. 4). We also observe interesting temporal transience in large exclosures, where infected questing nymph densities over time first have a rapid increase from their pre-exclosure equilibrium, followed by a long decrease to their post-exclosure equilibrium (Figs. 4*d* and 5*d*). This transient behavior is most significant with questing tick movement (Fig. 5*d*) and near the edge of the exclosure at the peak of the non-monotonic density gradient. Thus, it may be important in exclosure experiments to let the system equilibrate for a sufficient amount of time to capture its true nature.

## 4 Discussion

This work enhances the body of literature using mechanistic movement modeling to simulate TBP transmission and spread [46–50], adapting the approach to investigate how large mammal exclosures affect patterns in TBP risk over space. In doing so, we reconcile the conflicting results that have come from empirical studies using exclosures to determine how biodiversity loss affects TBP risk, from which smaller exclosures have tended to find a dilution effect while larger exclosures have found an amplification effect. We show that this scale dependence can result from a balance between a strong dilution effect at the exclosure’s edge, where the lack of deer increases infected tick densities, and a strong amplification effect far away from the exclosure’s edge, where the lack of deer decreases infected tick densities due to the limits of ticks’ rodent-based dispersal. This phenomenon, that dilution is dominant where deer have a strong influence while amplification is dominant where deer have a week influence, is consistent with the notion from previous empirical and theoretical work [25, 27, 28] that deer are amplification hosts at low densities and dilution hosts at high densities

Our results are also consistent with the one other study that has attempted a theoretical explanation of the scale-dependence of exclosure experiments [44]. However, their model of two homogeneous patches lacks the spatial nuance our model provides and is unable to explain spatial patterns of tick densities that have been empirically observed within exclosures. Averaging infected tick densities across the entire inside and outside of the exclosure and plotting for a range of exclosure sizes (Fig. 2), we recover the same pattern that finds, that the average infected tick density outside the exclosure stays fairly constant at an intermediate density for all exclosure sizes, while the average inside the exclosure is significantly higher than the outside for small exclosures, decreasing until it is lower than the outside for large exclosures. The shape of this decrease also matches that of [44], although their two-patch model parameterizes exclosure size by a “rodent flow rate” between the inside and outside patch which is inversely proportional to the actual exclosure size. By modeling space explicitly, we are able to parameterize exclosure size directly, rendering their model more easily interpretable. Our approach also allows one to simultaneously alter rodent movement rates independently of exclosure size (the “rodent flow rate” of [44] combines exclosure size and rodent movement into one parameter), incorporate movement from ticks in their free-living questing stage, and most importantly explore the risk gradients that exist within an exclosure.

The few empirical studies that have reported on tick gradients within exclosures have found that infected ticks were more abundant in the exclosure’s center than at its edge [14, 16, 31], in contrast to the results of our model without questing tick movement and the intuition that ticks should be nearly absent in the center of large exclosures due to the limits of rodent-based dispersal. However, our model is also able to recreate these reversed gradients once questing tick movement is taken into account. Specifically, these studies only investigated small exclosures (≤ 1 ha) [14, 31], for which our model with questing tick movement (Fig. 5*a*) reproduces the findings of [14, 31] that tick densities increased from the edges of their exclosures towards the centers. Though there is no similar data for larger exclosures, [16] later hypothesized that tick density gradients for large exclosures should be non-monotonic, first increasing and then decreasing eventually to zero going from the edge to center, as our model is also able to show (Fig. 5*b*). [31] and [16] justify their finding of low tick densities near the inside edge of the exclosure with their “tick sharing” hypothesis, which posits that rodents and deer sharing home ranges near the exclosure boundary can result in ticks being dropped off by rodents inside the exclosure, moving outside of the exclosure while questing, and then being picked up by deer outside. Though we do not explicitly model hosts’ home ranges, our analytic finding that nymph-deer interactions drive the reversed or non-monotonic gradients resulting from our model with questing tick movement gives theoretical support to the tick sharing mechanism.

The tick-borne disease risk gradients our model reveals are crucial to understanding the mechanisms driving scale dependence in exclosure experiment results. The maximum infected tick density within the exclosure near its edge indicates the maximum potential strength of the dilution effect, while the rate at which the gradient declines towards the center reveals how limited tick dispersal is by rodent movement. To make the model more applicable to interpreting results from exclosure experiments, we provide a formula for the gradient’s rate of exponential decline as a function of rodent dispersal rate and the average time ticks feed on rodents. We encourage future exclosure experiments to measure infected tick densities at points of varying distances from the exclosure’s edge, rather than averaging measurements across the entire exclosure. This can be accomplished by quantifying tick densities along transects that run parallel to the exclosure’s edge, recording tick densities and distance from the edge for each transect. Plotting the resulting infected tick density gradients can help disentangle the mechanisms driving these results, revealing the relative contributions of rodent-based and questing tick dispersal, tick sharing between rodents and large mammals, dilution, and amplification.

With our simple mechanistic modeling approach to capture tick, host and TBP life cycles and transmission, we ignore many important nuances common to the ecology and transmission of TBP systems that bear mentioning. Seasonality, for one, is a prominent factor of many tick life cycles with significant consequences for the spread of TBPs such as the vectors of Lyme disease and tick-borne encephalitis [54, 55]. And by considering only deer and rodent hosts, we omit a whole range of intermediate-sized hosts that may be either incompetent hosts able to enter the exclosure, or competent hosts unable to enter the exclosure [22]. The parameterization we present here only represents a narrow range of possibilities best meant to represent the mechanisms influencing exclosure results. However, some ticks may not feed on small mammals at any stage in their life, which could remove any significant patterns inside of the exclosure, as evidenced by the case of *Rhipicephalus pulchellus* from the data of Titcomb et al. [14, 16].

The host and tick movement component of our model also greatly simplifies the actual mobility patterns of these animals. No organisms move in perfectly random, undirected Brownian motions, as they react to heterogeneities in the landscape and countless other stimuli, while most mammals have home ranges in which they spend most of their time. We assumed that tick larvae never have any movement as this greatly simplifies the analytic results, but larvae have been observed to occasionally move small distances from their spawning site [56, 57]. Small mammal movement could be reduced if an individual is significantly burdened by ticks feeding on it. And tick or mammal movement could vary significantly between the inside and outside of the exclosure, as the loss of large herbivores inside the exclosure could yield higher plant densities. Other factors inside the exclosure, like lack of competition or predation, could be favorable for small mammals and increase their densities inside [14, 15].

On the other hand, our model with its 20 compartments varying over space could be considered too complicated to make any broad generalizations. Though we decided to stay faithful to the original processes modeled by [25, 44], our analytic results are generalizable enough to encompass a range of scenarios, including two-host ticks, non-linear transmission, and the absence of co-feeding transmission. Similarly, though we choose a circular exclosure shape for ease of simulation, our analytic results on tick density gradients can apply to any exclosure shape when it is sufficiently large. Our analytic results also rely on the simplifying assumptions that adult ticks do not feed on rodents and that rodents do not suffer any infection-based mortality, assumptions that are empirically supported and often made in the literature [25, 31, 53].

In spite of these limitations, our modeling approach is the first to reconcile the contrasting results of exclosure experiments while properly acknowledging the spatial heterogeneity that produces these contrasting results. Measuring these spatial gradients in tick densities and comparing them to our analytic formulae can elucidate their causal mechanisms and the nuances that drive observed relationships between biodiversity and tick-borne disease risk.

## Acknowledgements

We would like to thank Richard Ostfeld and Denis Patterson for helpful foundational discussions. A.S.F. and S.A.L. were supported by the NSF grant CCF1917819. Funding for S.A.L. was provided by the NSF grant DMS-2327711. This project was supported by the Stanford University’s Department of Psychiatry and Behavioral Sciences Lyme Disease Seed Grant Program. G.A.D. was partially supported also by the NSF grant DEB2011179.

## Code availability

All code to run the model and produce the figures from this paper can be found at https://github.com/freedmanari/TBP_exclosures.

