## Supplement for "A unifying theoretical framework for tick-borne disease risk to explain conflicting results of exclosure experiments across scales"

### 1 Model overview

Our model and parameters are adapted from Pugliese and Rosa's [1] model of tick-borne encephalitis in the presence of rodents (competent small mammal hosts), deer (incompetent large mammal hosts), and an exclosure that keeps out the deer. While their model has two homogeneous patches—an inside-exclosure patch and an outside-exclosure patch—with a parameter controlling the flow of rodents between the two patches which is roughly inversely proportional to the exclosure's size, we bring their model explicitly into two-dimensional space. Doing so allows us to forgo the assumption that everything is homogeneous inside or outside the exclosure, and parameterize exclosure size directly. While one can use this model to simulate an exclosure of any shape, we choose a circular exclosure because this allows us to assume everything is radially symmetric centered around the center of the exclosure, effectively reducing the model to one dimension for simulation purposes.

We simulate the following partial differential equations (PDEs) in R using the package `ReacTran` [2], approximating continuous space in polar coordinates by divide the $r$-axis from 0–1,000 m into 800 slices. The exclosure is a circular area centered at the same center as the larger 1,000 m-radius circular domain we are modeling over, with the exclosure radius ranging up to a maximum of 300 m. We first let the model equilibrate over 3,000 days assuming there is no space or exclosure; we then plug in the equilibrium values as initial conditions for the model with space, which we also let equilibrate over 3,000 days to get the spatially-explicit exclosure model's equilibrium. Doing so simulates what would happen in an exclosure was suddenly placed in an ecosystem that was previously at an exclosure-free equilibrium. The spatially-explicit model's initial conditions are constant over space for all state variables, except the state variables for deer or ticks feeding on deer, whose initial conditions are 0 inside the exclosure and constant outside the exclosure at their pre-space equilibria.

The only change we make from Pugliese and Rosa's base model is to allow co-feeding transmission to happen from infected adult ticks, as well as from infected nymph ticks (this is how it is described in their main text, but the actual equations in their model only include co-feeding transmission from nymphs). In the parameterization we present in this paper, we assume that the rate of co-feeding transmission from adults, $\lambda^A$, is equal to their rate of co-feeding transmission from nymphs, $\lambda^N$.

Our model also allows us to parameterize rodent movement independently from exclosure size, using the rodent's diffusion coefficient $D_1$. We parameterize rodent movement from the more easily measured quantity of the rodent's average daily movement rate $v_1$ (the average distance a rodent would travel in a day given it is moving under Brownian motion). Since the distance a 2-dimensional Brownian motion travels after time $\tau$ with diffusion coefficient $D$ is distributed as a Rayleigh distribution [3] with scale parameter $\sqrt{2D\tau}$, the average of this distribution with $\tau = 1$ day gives $v_1 = \sqrt{\pi D_1}$. Equivalently, $D_1 = \frac{v_1^2}{\pi}$. Our model also gives us the opportunity to explicitly model deer movement (we assume here that deer birth and death rates are equal so that the deer population is constant across space and time, but their movement may still have an impact on tick dispersal), as well as questing tick movement in all of the tick life stages. For these, we also parameterize movement by calculating their respective diffusion coefficients from their assumed average daily movement rates, in the same way we do for rodents.

All state variables and parameter values, are shown in the tables just below. All parameter values come from [1] except animal movement rates—which we vary over an assumed range—and we multiply their mammal-tick contact rates by a factor of 2, 3, or 5 depending on the scenario we are modeling (most of our simulations assume the factor of 3, and no outbreaks occur if we do not multiply the contact rates by some factor greater than 1). In the model equations, $\dot{X} = \frac{\partial X}{\partial t}$ is the time derivative of variable $X$, and $\Delta = \frac{\partial^2}{\partial x^2} + \frac{\partial^2}{\partial y^2}$ is the Laplacian operator, where $x$ and $y$ are the Cartesian coordinates of the two-dimensional space. And $\Delta^{\text{out}}$ is the Laplacian operator restricted to the outside of the exclosure, for deer and ticks feeding on deer, equal to the normal Laplacian outside the exclosure but equal to 0 inside the exclosure.

### 2 Tables of symbols and parameter values

| Symbol | Description |
|---|---|
| $H_1/H_2$ | Total rodent/deer density |
| $T_{F_1}/T_{F_2}$ | Total density of ticks feeding on rodents/deer |
| $L/N/A$ | Density of tick larvae/nymphs/adults |
| $X_Q$ | Density of questing ticks in life stage $X$ |
| $X_{F_1}/X_{F_2}$ | Density of ticks of type $X$ feeding on rodents/deer |
| $X^s/X^i$ | Density of rodents/ticks of type $X$ which are susceptible/infected |
| $X^e$ | Density of feeding ticks of type $X$ which are exposed (but not infectious) |
| $H_1^r$ | Density of rodents which are susceptible |

Table 1: Model state variables, all with unit (individuals $\mathrm{m}^{-2}$)

| Symbol | Description | Value and unit |
|---|---|---|
| $b_1/b_2$ | Rodent/deer birth rate | .00821/0 $\mathrm{day}^{-1}$ |
| $d_1/d_2$ | Rodent/deer death rate | .0037/0 $\mathrm{day}^{-1}$ |
| $K_1/K_2$ | Rodent/deer carrying capacity | .0015/.00001 individuals $\mathrm{m}^{-2}$ |
| $\gamma$ | Rodent recovery rate | .3 $\mathrm{day}^{-1}$ |
| $\alpha$ | Rodent infection-induced mortality rate | .33 $\mathrm{day}^{-1}$ |
| $b^T$ | Average egg production per fed adult tick | 2,000 |
| $c$ | Density-dependent death rate of ticks | 2,500 $\mathrm{m}^2$ $\mathrm{ticks}^{-1}$ |
| $d^X$ | Death rate of ticks of type $X(=L/N/A)$ | .0365/.015/.00625 $\mathrm{day}^{-1}$ |
| $\sigma^X$ | Detachment rate of ticks of type $X(=L/N/A)$ | .28/.22/.12 $\mathrm{day}^{-1}$ |
| $m^X$ | Molting success probability for ticks of type $X(=L/N)$ | .15/.15 |
| $\beta_1^X$ | Contact rate between questing ticks of type $X(=L/N/A)$ and rodents | $\omega \cdot$ (150/5/.2) $\mathrm{individuals}^{-1}$ $\mathrm{day}^{-1}$ |
| $\beta_2^X$ | Contact rate between questing ticks of type $X(=L/N/A)$ and deer | $\omega \cdot$ (800/2,500/2,500) $\mathrm{individuals}^{-1}$ $\mathrm{day}^{-1}$ |
| $\omega$ | Contact rate multiplier for low/medium/high contact scenarios | 2/3/5 |
| $\lambda^X$ | Probability of getting infected via co-feeding from tick of type $X(=N/A)$ | .55/.55 |
| $v_1$ | Average daily movement for rodents in lower/medium/high movement scenarios | 10/25/50 m $\mathrm{day}^{-1}$ |
| $v_2$ | Average daily movement for deer | 250 m $\mathrm{day}^{-1}$ |
| $v_X$ | Average daily movement for ticks of type $X(=L/N/A)$, *when there's tick movement* | 0/5/5 m $\mathrm{day}^{-1}$ |
| $D_1/D_2$ | Diffusion coefficient for rodents/deer, equal to $\frac{v_1^2}{\pi}$ / $\frac{v_2^2}{\pi}$ | $\mathrm{m}^2$ $\mathrm{day}^{-1}$ |
| $D^X$ | Diffusion coefficient for ticks of type $X(=L/N/A)$, equal to $\frac{v_X^2}{\pi}$ | $\mathrm{m}^2$ $\mathrm{day}^{-1}$ |

Table 2: Model parameters and other symbols, with their values and units

### 3 PDEs with spatial dynamics

#### 3.1 Host dynamics

$$\mu_1(H_1) = d_1 + \tfrac{H_1}{K_1}(b_1 - d_1) \tag{1}$$

$$\mu_2(H_2) = d_2 + \tfrac{H_2}{K_2}(b_2 - d_2) \tag{2}$$

$$\dot{H}_1^s = b_1 H_1 - \left(\beta_1^N N_Q^i + \beta_1^A A_Q^i\right) H_1^s - \mu_1(H_1) H_1^s + D_1 \Delta H_1^s \tag{3}$$

$$\dot{H}_1^i = \left(\beta_1^N N_Q^i + \beta_1^A A_Q^i\right) H_1^s - \left(\gamma + \mu_1(H_1) + \alpha\right) H_1^i + D_1 \Delta H_1^i \tag{4}$$

$$\dot{H}_1^r = \gamma H_1^i - \mu_1(H_1) H_1^r + D_1 \Delta H_1^r \tag{5}$$

$$\dot{H}_2 = b_2 H_2 - \mu_2(H_2) H_2 + D_2 \Delta^{\mathrm{out}} H_2 \tag{6}$$

$$\tag{7}$$

##### 3.2 Tick dynamics

###### 3.2.1 Larvae

$$\dot{L}_Q = \frac{b^T}{1+cT_{F_1}}\sigma^A A_{F_1} + \frac{b^T}{1+cT_{F_2}}\sigma^A A_{F_2} - \left(\beta_1^L H_1 + \beta_2^L H_2\right) L_Q - d^L L_Q + D^L \Delta L_Q \tag{8}$$

$$\dot{L}^s_{F_1} = \exp\left(-\frac{\lambda^N N^i_{F_1} + \lambda^A A^i_{F_1}}{H_1}\right) \beta_1^L \left(H_1^s + H_1^r\right) L_Q - \sigma^L L^s_{F_1} + D_1 \Delta L^s_{F_1} \tag{9}$$

$$\dot{L}^e_{F_1} = \beta_1^L H_1^i L_Q + \left[1 - \exp\left(-\frac{\lambda^N N^i_{F_1} + \lambda^A A^i_{F_1}}{H_1}\right)\right] \beta_1^L \left(H_1^s + H_1^r\right) L_Q - \sigma^L L^e_{F_1} + D_1 \Delta L^e_{F_1} \tag{10}$$

$$\dot{L}^s_{F_2} = \beta_2^L H_2 L_Q - \sigma^L L^s_{F_2} + D_2 \Delta^{\text{out}} L^s_{F_2} \tag{11}$$

###### 3.2.2 Nymphs

$$\dot{N}^s_Q = m^L \sigma^L \left(L^s_{F_1} + L^s_{F_2}\right) - \left(\beta_1^N H_1 + \beta_2^N H_2\right) N^s_Q - d^N N^s_Q + D^N \Delta N^s_Q \tag{12}$$

$$\dot{N}^i_Q = m^L \sigma^L L^e_{F_1} - \left(\beta_1^N H_1 + \beta_2^N H_2\right) N^i_Q - d^N N^i_Q + D^N \Delta N^i_Q \tag{13}$$

$$\dot{N}^s_{F_1} = \exp\left(-\frac{\lambda^N N^i_{F_1} + \lambda^A A^i_{F_1}}{H_1}\right) \beta_1^N \left(H_1^s + H_1^r\right) N^s_Q - \sigma^N N^s_{F_1} + D_1 \Delta N^s_{F_1} \tag{14}$$

$$\dot{N}^e_{F_1} = \beta_1^N H_1^i N^s_Q + \left[1 - \exp\left(-\frac{\lambda^N N^i_{F_1} + \lambda^A A^i_{F_1}}{H_1}\right)\right] \beta_1^N \left(H_1^s + H_1^r\right) N^s_Q - \sigma^N N^e_{F_1} + D_1 \Delta N^e_{F_1} \tag{15}$$

$$\dot{N}^i_{F_1} = \beta_1^N H_1 N^i_Q - \sigma^N N^i_{F_1} + D_1 \Delta N^i_{F_1} \tag{16}$$

$$\dot{N}^s_{F_2} = \beta_2^N H_2 N^s_Q - \sigma^N N^s_{F_2} + D_2 \Delta^{\text{out}} N^s_{F_2} \tag{17}$$

$$\dot{N}^i_{F_2} = \beta_2^N H_2 N^i_Q - \sigma^N N^i_{F_2} + D_2 \Delta^{\text{out}} N^i_{F_2} \tag{18}$$

###### 3.2.3 Adults

$$\dot{A}^s_Q = m^N \sigma^N \left(N^s_{F_1} + N^s_{F_2}\right) - \left(\beta_1^A H_1 + \beta_2^A H_2\right) A^s_Q - d^A A^s_Q + D^A \Delta A^s_Q \tag{19}$$

$$\dot{A}^i_Q = m^N \sigma^N \left(N^e_{F_1} + N^i_{F_1} + N^i_{F_2}\right) - \left(\beta_1^A H_1 + \beta_2^A H_2\right) A^i_Q - d^A A^i_Q + D^A \Delta A^i_Q \tag{20}$$

$$\dot{A}^s_{F_1} = \beta_1^A H_1 A^s_Q - \sigma^A A^s_{F_1} + D_1 \Delta A^s_{F_1} \tag{21}$$

$$\dot{A}^i_{F_1} = \beta_1^A H_1 A^i_Q - \sigma^A A^i_{F_1} + D_1 \Delta A^i_{F_1} \tag{22}$$

$$\dot{A}_{F_2} = \beta_2^A H_2 \left(A^s_Q + A^i_Q\right) - \sigma^A A_{F_2} + D_2 \Delta^{\text{out}} A_{F_2} \tag{23}$$

### 4 Analytic results

In order to make the model (1)–(23) more tractable, we consider the model over the entire domain $\mathbb{R}^2$, with the exclosure represented by the area $\{(x, y) \in \mathbb{R}^2 \mid x \geq 0\}$ with non-negative $x$-coordinates. This approximation works well for large circular exclosures. Furthermore, we consider initial conditions that are constant along the $y$-axis, i.e. $X(x, y, t) = X(x, 0, t)\ \forall y \in \mathbb{R}$ for any state variable $X$. We can thus reduce the model to one dimension along just the $x$-axis, and notate each state variable $X$ over space and time as $X(x, t)$. We notate the equilibrium value of state variable $X$ at point in space $x$ as $\overline{X}(x)$. We seek to characterize the equilibrium infected nymph density at all points inside the exclosure, $\overline{N^i_q}(x)$ for $x \geq 0$.

Lastly, we make the simplifications that $\beta_1^A = 0$ (i.e. adult ticks do not feed on rodents) and that $\alpha = 0$ (i.e. rodents do not suffer any infection-based mortality). Both of these assumptions have been found to be supported and are commonly taken to be true [4–6]. Furthermore, we start out with $H_1(x, 0) = K_1$, making it so that the total rodent densities are constant over space and time, i.e. $H_1(x, t) = H_1^s(x, t) + H_1^i(x, t) + H_1^r(x, t) = K_1\ \forall x, t$. Since deer already don't suffer any infection-based mortality and we also start them out at their carrying capacity $H_2(x, 0) = K_2$ for $x < 0$, it is also true that deer densities are constant over space and time outside the exclosure, i.e. $H_2(x, t) = K_2\ \forall x < 0, t$.

First we provide the following lemma:

**Lemma 1.**
$$\int_0^\infty \frac{1}{\sqrt{\pi\tau}} e^{-\frac{x^2}{4D\tau}-\delta\tau} d\tau = \frac{1}{\sqrt{\delta}} e^{-\sqrt{\frac{\delta}{D}}|x|} \tag{24}$$

*for all $x \in \mathbb{R}$ and $D, \delta > 0$*

*Proof.* First, we see that

$$\begin{aligned}
&\frac{d}{d\tau}\left(\int_0^{\sqrt{\frac{x^2}{4D\tau}}+\sqrt{\delta\tau}} e^{-z^2+\sqrt{\frac{\delta}{D}}|x|}dz - \int_0^{\sqrt{\frac{x^2}{4D\tau}}-\sqrt{\delta\tau}} e^{-z^2-\sqrt{\frac{\delta}{D}}|x|}dz\right)\\
&= \frac{1}{2}\left(-\sqrt{\frac{x^2}{4D\tau^3}}+\sqrt{\frac{\delta}{\tau}}\right) e^{-\left(\sqrt{\frac{x^2}{4D\tau}}+\sqrt{\delta\tau}\right)^2+\sqrt{\frac{\delta}{D}}|x|} + \frac{1}{2}\left(\sqrt{\frac{x^2}{4D\tau^3}}+\sqrt{\frac{\delta}{\tau}}\right) e^{-\left(\sqrt{\frac{x^2}{4D\tau}}-\sqrt{\delta\tau}\right)^2-\sqrt{\frac{\delta}{D}}|x|}\\
&= \sqrt{\frac{\delta}{\tau}} e^{-\frac{x^2}{4D\tau}-\delta\tau},
\end{aligned}$$

and so

$$\int_{t_1}^{t_2} \frac{1}{\sqrt{\pi\tau}} e^{-\frac{x^2}{4D\tau}-\delta\tau} d\tau = \frac{1}{2\sqrt{\delta}}\left[e^{\sqrt{\frac{\delta}{D}}|x|}\operatorname{erf}\left(\sqrt{\frac{x^2}{4D\tau}}+\sqrt{\delta\tau}\right) - e^{-\sqrt{\frac{\delta}{D}}|x|}\operatorname{erf}\left(\sqrt{\frac{x^2}{4D\tau}}-\sqrt{\delta\tau}\right)\right]_{\tau=t_1}^{t_2}$$

for any $t_1, t_2 > 0$, where erf is the error function, defined as $\operatorname{erf}(z) = \frac{2}{\sqrt{\pi}}\int_0^z e^{-t^2}dt$. As we take the limits $t_1 \to 0^+$ and $t_2 \to \infty$, using the fact that $\lim_{z\to\pm\infty} \operatorname{erf}(z) = \pm 1$, we get the desired result. □

#### 4.1 Gradient of $\overline{N_Q^i}$ without questing tick movement

Assuming there is no questing tick movement in the larval or nymph stages ($D^L = D^N = 0$), we are able to prove the following result, that the infected questing nymph density gradient inside the exclosure exponentially declines at rate $\sqrt{\frac{\sigma^L}{D_1}}$ as we move away from the edge of the exclosure:

**Theorem 1.** *If $\beta_1^A = 0$, $\alpha = 0$, and $D^L = D^N = 0$, then*

$$\overline{N_Q^i}(x) = \frac{m^L\sigma^L}{\beta_1^N K_1 + d^N}\overline{L_{F_1}^e}(0^+)e^{-\sqrt{\frac{\sigma^L}{D_1}}x} \quad \forall x \geq 0, \tag{25}$$

*where*

$$\overline{L_{F_1}^e}(0^+) = \lim_{x\to 0+}\overline{L_{F_1}^e}(x) = \int_{-\infty}^0 \frac{\overline{\varphi}(y)}{\sqrt{4D_1\sigma^L}} e^{\sqrt{\frac{\sigma^L}{D_1}}y}dy \tag{26}$$

*and*

$$\overline{\varphi}(x) = \beta_1^L\left[\overline{H_1^i}(x) + \left(1-\exp\left(-\frac{\lambda^N\overline{N_{F_1}^i}(x)+\lambda^A\overline{A_{F_1}^i}(x)}{K_1}\right)\right)\left(K_1-\overline{H_1^i}(x)\right)\right]\overline{L_Q}(x). \tag{27}$$

*Consequently, $\overline{N_Q^i(x)}$ has an exponential decay rate of $\sqrt{\frac{\sigma^L}{D_1}}$ for all $x \geq 0$.*

*Proof.* With $\beta_1^A = 0$, there is never any influx into $A_{F_1}^s$ or $A_{F_1}^i$ by (21)–(22) (disregarding the diffusion terms which will approach 0), and so $\overline{A_{F_1}^s}(x) = \overline{A_{F_1}^i}(x) = 0\ \forall x \in \mathbb{R}$. Similarly, since $H_2(x) = 0$ for all non-negative $x$, there is no influx into $A_{F_2}$ at all for any non-negative $x$ by (23), and so $\overline{A_{F_2}}(x) = 0\ \forall x \geq 0$. Thus, at equilibrium there will be no influx into $L_Q(x)$ for any non-negative $x$ by (8), so $\overline{L_Q}(x) = 0\ \forall x \geq 0$.

With $D^N = 0$, and $\alpha = 0$ rendering $H_1(x,t) = K_1$ a constant for all $x$ and $t$, (13) gives us

$$\overline{N_Q^i}(x) = \frac{m^L\sigma^L}{\beta_1 K_1 + d^N}\overline{L_{F_1}^e}(x)\ \forall x \geq 0, \tag{28}$$

since $H_2(x) = 0$ for non-negative $x$.

Now every tick in $L^e_{F_1}(x,t)$ can be characterized by the location $y$ at which they first flowed (or were initialized) into the class, and the amount of time $\tau \leq t$ that they have been in the class for. Since $\alpha = 0$ means $H^s_1 + H^r_1 = K_1 - H^i_1$, (10) gives that the new ticks flowing into $L^e_{F_1}$ at location $x$ and time $t$ is

$$\varphi(x,t) = \beta^L_1 \left[ H^i_1(x,t) + \left(1 - \exp\left(-\frac{\lambda^N N^i_{F_1}(x,t) + \lambda^A A^i_{F_1}(x,t)}{K_1}\right)\right)\left(K_1 - H^i_1(x,t)\right)\right] L_Q(x,t). \tag{29}$$

In general, for a point source starting at position $x$ with density $\varphi$ and diffusing out with diffusion coefficient $D$, the density time $\tau$ later and at position $y$ is given by $\frac{\varphi}{\sqrt{4\pi D\tau}} e^{-\frac{(x-y)^2}{4D\tau}}$ [7]. Lastly, ticks in $L^e_{F_1}$ leave the class at rate $\sigma^L$, so the probability that a tick stays in the class for at least time $\tau$ is defined by $e^{-\sigma^L \tau}$. All together, this gives $L^e_{F_1}(x,t) = \int_{-\infty}^{\infty}\int_0^t \frac{\varphi(y,t-\tau)}{\sqrt{4\pi D_1 \tau}} e^{-\frac{(x-y)^2}{4D\tau}} e^{-\sigma^L \tau} d\tau dy + \int_{-\infty}^{\infty} \frac{\varphi(y,0)}{\sqrt{4\pi D_1 t}} e^{-\frac{(x-y)^2}{4Dt}} e^{-\sigma^L t} dy$. Going to equilibrium as $t \to \infty$, the second term here will vanish, and $\varphi(x,t)$ will tend to $\overline{\varphi}(x)$ as given by (27) with $\overline{\varphi}(x) = 0 \ \forall x \geq 0$, leaving

$$\overline{L^e_{F_1}}(x) = \int_{-\infty}^{0}\int_0^{\infty} \frac{\overline{\varphi}(y)}{\sqrt{4\pi D_1 \tau}} e^{-\frac{(x-y)^2}{4D_1\tau}} e^{-\sigma^L \tau} d\tau dy \quad \forall x \in \mathbb{R}. \tag{30}$$

Using Lemma 1, we can simplify

$$\overline{L^e_{F_1}}(x) = \int_{-\infty}^{0} \frac{\overline{\varphi}(y)}{\sqrt{4 D_1 \sigma^L}} e^{-\sqrt{\frac{\sigma^L}{D_1}}|x-y|} dy \quad \forall x \in \mathbb{R}. \tag{31}$$

For $x \geq 0$, this gives us

$$\overline{L^e_{F_1}}(x) = \overline{L^e_{F_1}}(0^+) e^{-\sqrt{\frac{\sigma^L}{D_1}}x} \tag{32}$$

by (26), from which (28) give us the desired result.

□

**Remark.** Note that even when $\alpha \neq 0$, we can still get the expression

$$\overline{N^i_Q}(x) = \frac{m^L \sigma^L}{\beta^N_1 \overline{H_1}(x) + d^N} \overline{L^e_{F_1}}(0^+) e^{-\sqrt{\frac{\sigma^L}{D_1}}x} \quad \forall x \geq 0 \tag{33}$$

for $\overline{N^i_Q}(x)$. Though $\overline{H_1}(x)$ need not be uniform inside the exclosure when $\alpha > 0$, the differences in $\overline{H_1}(x)$ within the exclosure will likely be relatively minor for any realistic values of $\alpha$, and thus not disturb the monotonicity of the $\overline{N^i_Q}(x)$ gradient inside the exclosure.

#### 4.2 Gradient of $\overline{N^i_Q}$ with questing tick movement

Now we investigate what happens to the $\overline{N^i_Q}(x)$ gradient inside the exclosure when we allow questing tick movement in nymphs, i.e. $D^N > 0$ but we still enforce $D^L = 0$ (as larvae have relatively little movement while questing compared to later stages). We prove that when $\beta^N_2$, the contact rate between nymphs and deer, is 0, the infected questing nymph gradient inside the exclosure is of the form $\overline{N^i_Q}(x) = a_1 e^{-r_1 x} - a_2 e^{-r_2 x} \ \forall x \geq 0$ and monotonically decreasing. We thus show that the non-monotonic gradients we observe within the exclosures in the main text are solely the result of high contact rates between nymphs and deer outside the exclosure, as questing ticks inside the exclosure that are closer to the boundary are more likely to get picked up by a mammal host in a short amount of time due to their proximity to deer.

We first provide the following lemma:

**Lemma 2.** *Any function $f$ of the form $f(x) = a_1 e^{-r_1 x} - a_2 e^{-r_2 x} \ \forall x \geq 0$ with $a_1, r_1, a_2, r_2 > 0$ that satisfies $f(x) > 0 \ \forall x \geq 0$ will be monotically decreasing for all $x \geq 0$ if and only if $a_1 r_1 \geq a_2 r_2$. Furthermore, if $a_1 r_1 < a_2 r_2$, then $f(x)$ is non-monotonic for $x \geq 0$, with an initial increase followed by a decline to 0. In both cases, $f(x)$ as $x \to \infty$ has an the asymptotic exponential decay rate of $r_1$, i.e. $\lim_{x\to\infty} \frac{d}{dx} \log f(x) = -r_1$.*

*Proof.* We must have $r_1 \le r_2$, since otherwise $a_2 e^{-r_2 x}$ would dominate $a_1 e^{-r_1 x}$ eventually and we would have $f(x) < 0$ for sufficiently large $x$. If $r_1 = r_2$, then $f(x) = (a_1 - a_2)e^{-r_1 x}$, so it must be true that $a_1 > a_2$ to ensure $f(x) > 0$. Thus, this case gives us $a_1 r_1 > a_2 r_2$ and $f(x)$ monotonically decreasing at an exponential decay at rate $r_1$, so the lemma is satisfied for $r_1 = r_2$.

Otherwise, $r_1 < r_2$. Now we see that $f'(x) = -a_1 r_1 e^{-r_1 x} + a_2 r_2 e^{-r_2 x} \le 0\ \forall x \ge 0$ if $a_1 r_1 \ge a_2 r_2$, since $r_1 - r_2 < 0$ and $\frac{a_2 r_2}{a_1 r_1} \le 1$ ensure that $\frac{a_2 r_2}{a_1 r_1} e^{(r_1 - r_2)x} \le 1\ \forall x \ge 0$. Thus, $f(x)$ will be monotonically decreasing for $x \ge 0$. And if $a_1 r_1 < a_2 r_2$, then $f'(0) = -a_1 r_1 + a_2 r_2 > 0$, giving $f$ an initial increase, followed by a decline to 0 for larger $x$ as $r_1 < r_2$ ensures that $e^{-r_1 x}$ will eventually come to dominate $e^{-r_2 x}$, leading $f(x)$ to be non-monotonic over $x \ge 0$. No matter the case, $r_1 < r_2$ ensures $e^{-r_1 x}$ will eventually dominate $e^{-r_2 x}$, leading to $f$ having an asymptotic exponential decay rate of $r_1$. □

**Theorem 2.** *If $\beta_1^A = \beta_2^N = 0$, $\alpha = 0$, and $D^L = 0$ but $D^N > 0$, then*

$$\overline{N_Q^i}(x) = \frac{m^L \sigma^L}{\sqrt{4D^N(\beta_1^N K_1 + d^N)}} \left(\psi_1 e^{-\sqrt{\eta_1}x} - \psi_2 e^{-\sqrt{\eta_2}x}\right) \quad \forall x \ge 0, \tag{34}$$

*where*

$$\begin{aligned}
\eta_1 &= \frac{\sigma^L}{D_1} \\
\eta_2 &= \frac{\beta_1^N K_1 + d^N}{D^N} \\
\psi_1 &= \frac{2\sqrt{\eta_2}}{\eta_2 - \eta_1} \int_{-\infty}^0 \frac{\overline{\varphi}(y)}{\sqrt{4D_1\sigma^L}} e^{\sqrt{\eta_1}y} dy = \frac{2\sqrt{\eta_2}}{\eta_2 - \eta_1} \overline{L_{F_1}^e}(0^+) \\
\psi_2 &= \frac{2\sqrt{\eta_1}}{\eta_2 - \eta_1} \int_{-\infty}^0 \frac{\overline{\varphi}(y)}{\sqrt{4D_1\sigma^L}} e^{\sqrt{\eta_2}y} dy,
\end{aligned}$$

*with $\overline{L_{F_1}^e}(0^+)$ and $\overline{\varphi}(x)$ defined as in (26)–(27). Furthermore, $\overline{N_Q^i}(x)$ is monotonically decreasing over all $x \ge 0$.*

*Proof.* The same logic from the proof of Theorem 1 still holds, except (28) no longer holds with $D^N > 0$. Specifically, (30)–(32) are still valid. To calculate $\overline{N_Q^i}(x)$, we consider ticks in $N_Q^i$ entering the class at location $y$ at rate $m^L \sigma^L \overline{L_{F_1}^e}(y)$, diffusing with diffusion coefficient $D^N$, and leaving the class at rate $\beta_1^N K_1 + d^N$ both inside and outside the exclosure (since $\beta_2^N = 0$), giving us

$$\overline{N_Q^i}(x) = \int_{-\infty}^{\infty} \int_0^{\infty} \frac{m^L \sigma^L \overline{L_{F_1}^e}(y)}{\sqrt{4\pi D^N \tau}} e^{-\frac{(x-y)^2}{4D^N \tau}} e^{-(\beta_1^N K_1 + d^N)\tau} d\tau dy \quad \forall x \in \mathbb{R}. \tag{35}$$

Applying Lemma 1 and plugging in (32), we derive for $x \ge 0$

$$\begin{aligned}
\overline{N_Q^i}(x) &= \frac{m^L \sigma^L}{\sqrt{4D^N(\beta_1^N K_1 + d^N)}} \int_{-\infty}^{\infty} \overline{L_{F_1}^e}(y) e^{-\sqrt{\eta_2}|x-y|} dy \\
&= \frac{m^L \sigma^L}{\sqrt{4D^N(\beta_1^N K_1 + d^N)}} \Bigg[ \int_{-\infty}^0 \overline{L_{F_1}^e}(y) e^{-\sqrt{\eta_2}(x-y)} dy \\
&\quad + \int_0^x \overline{L_{F_1}^e}(0^+) e^{-\sqrt{\eta_1}y} e^{-\sqrt{\eta_2}(x-y)} dy + \int_x^{\infty} \overline{L_{F_1}^e}(0^+) e^{-\sqrt{\eta_1}y} e^{-\sqrt{\eta_2}(y-x)} dy \Bigg] \\
&= \frac{m^L \sigma^L}{\sqrt{4D^N(\beta_1^N K_1 + d^N)}} \Bigg[ \left( \int_{-\infty}^0 \overline{L_{F_1}^e}(y) e^{\sqrt{\eta_2}y} dy \right) e^{-\sqrt{\eta_2}x} \\
&\quad + \frac{\overline{L_{F_1}^e}(0^+)}{\sqrt{\eta_2} - \sqrt{\eta_1}} \left( e^{-\sqrt{\eta_1}x} - e^{-\sqrt{\eta_2}x} \right) + \frac{\overline{L_{F_1}^e}(0^+)}{\sqrt{\eta_1} + \sqrt{\eta_2}} e^{-\sqrt{\eta_1}x} \Bigg].
\end{aligned} \tag{36}$$

For the integral in (36), we can plug in (31) and switch the order of integration to get

$$\begin{aligned}
\int_{-\infty}^{0} \overline{L^e_{F_1}}(y) e^{\sqrt{\eta_2} y} dy &= \int_{-\infty}^{0} \int_{-\infty}^{0} \frac{\overline{\varphi}(z)}{\sqrt{4D_1\sigma^L}} e^{-\sqrt{\eta_1}|y-z|} e^{\sqrt{\eta_2} y} dz\, dy \\
&= \int_{-\infty}^{0} \int_{z}^{0} \frac{\overline{\varphi}(z)}{\sqrt{4D_1\sigma^L}} e^{-\sqrt{\eta_1}(y-z)} e^{\sqrt{\eta_2} y} dy\, dz \\
&+ \int_{-\infty}^{0} \int_{-\infty}^{z} \frac{\overline{\varphi}(z)}{\sqrt{4D_1\sigma^L}} e^{-\sqrt{\eta_1}(z-y)} e^{\sqrt{\eta_2} y} dy\, dz \\
&= \frac{1}{\sqrt{\eta_2}-\sqrt{\eta_1}} \int_{-\infty}^{0} \frac{\overline{\varphi}(z)}{\sqrt{4D_1\sigma^L}} \left( e^{\sqrt{\eta_1} z} - e^{\sqrt{\eta_2} z} \right) dz \\
&+ \frac{1}{\sqrt{\eta_1}+\sqrt{\eta_2}} \int_{-\infty}^{0} \frac{\overline{\varphi}(z)}{\sqrt{4D_1\sigma^L}} e^{\sqrt{\eta_2} z} dz \\
&= \frac{\overline{L^e_{F_1}}(0^+)}{\sqrt{\eta_2}-\sqrt{\eta_1}} - \frac{2\sqrt{\eta_1}}{\eta_2-\eta_1} \int_{-\infty}^{0} \frac{\overline{\varphi}(y)}{\sqrt{4D_1\sigma^L}} e^{\sqrt{\eta_2} y} dy, \qquad (37)
\end{aligned}$$

where we use (26) in the last step. Plugging (37) into (36) and simplifying yields the result (34).

Now to show that $\overline{N^i_Q}(x)$ is monotonically decreasing for all $x \geq 0$, we examine two cases and show that Lemma 2 holds for both of them. Since $\overline{N^i_Q}(x) > 0$ everywhere (unless the disease dies out, then $\overline{N^i_Q}(x) = 0$ everywhere and is monotonically decreasing trivially), it suffices to choose $a_1, r_1, a_2, r_2 > 0$ such that $\overline{N^i_Q}(x) = a_1 e^{-r_1 x} - a_2 e^{-r_2 x}\ \forall x \geq 0$ and show that $a_1 r_1 > a_2 r_2$ in each case.

In the first case, when $\eta_1 < \eta_2$, we have $\psi_1, \psi_2 > 0$, so we choose $a_1 = \frac{m^L\sigma^L}{\sqrt{4D^N(\beta_1^N K_1 + d^N)}}\psi_1$, $r_1 = \sqrt{\eta_1}$, $a_2 = \frac{m^L\sigma^L}{\sqrt{4D^N(\beta_1^N K_1 + d^N)}}\psi_2$, and $r_2 = \sqrt{\eta_2}$. Then $a_1 r_1 > a_2 r_2$ simplifies to $\int_{-\infty}^{0} \frac{\overline{\varphi}(y)}{\sqrt{4D_1\sigma^L}} e^{\sqrt{\eta_1} y} dy > \int_{-\infty}^{0} \frac{\overline{\varphi}(y)}{\sqrt{4D_1\sigma^L}} e^{\sqrt{\eta_2} y} dy$, which is true by $\eta_1 < \eta_2$, so Lemma 2 is satisfied.

In the second case, when $\eta_1 > \eta_2$, we have $\psi_1, \psi_2 < 0$, so we choose $a_1 = -\frac{m^L\sigma^L}{\sqrt{4D^N(\beta_1^N K_1 + d^N)}}\psi_2$, $r_1 = \sqrt{\eta_2}$, $a_2 = -\frac{m^L\sigma^L}{\sqrt{4D^N(\beta_1^N K_1 + d^N)}}\psi_1$, and $r_2 = \sqrt{\eta_1}$. Then we see that $a_1 r_1 > a_2 r_2$ and thus Lemma 2 are still satisfied, since now $\int_{-\infty}^{0} \frac{\overline{\varphi}(y)}{\sqrt{4D_1\sigma^L}} e^{\sqrt{\eta_2} y} dy > \int_{-\infty}^{0} \frac{\overline{\varphi}(y)}{\sqrt{4D_1\sigma^L}} e^{\sqrt{\eta_1} y} dy$.

Here we disregard the case of $\eta_1 = \eta_2$, as in practice this will never occur exactly, and this leads to a different functional form for $\overline{N^i_Q}(x)$ which is also monotonically decreasing for $x \geq 0$. We briefly address this in the following remark.

□

**Remark.** In the case of $\eta_1 = \eta_2$, using L'Hôpital's rule in the limit of $\zeta = \eta_2 - \eta_1 \to 0$ on (34) for $x \geq 0$ yields

$$\begin{aligned}
\overline{N^i_Q}(x) &= \frac{m^L\sigma^L}{\sqrt{4D^N(\beta_1^N K_1 + d^N)}} \lim_{\zeta\to 0} \frac{2\sqrt{\eta_1+\zeta}\,\overline{L^e_{F_1}}(0^+) e^{-\sqrt{\eta_1}x} - 2\sqrt{\eta_1}\int_{-\infty}^{0} \frac{\overline{\varphi}(y)}{\sqrt{4D_1\sigma^L}} e^{\sqrt{\eta_1+\zeta}(y-x)} dy}{\zeta} \\
&= \frac{m^L\sigma^L}{\sqrt{4D^N(\beta_1^N K_1 + d^N)}} \left( \left( \frac{1}{\sqrt{\eta_1}} + x \right) \overline{L^e_{F_1}}(0^+) + \int_{-\infty}^{0} \frac{-y\overline{\varphi}(y)}{\sqrt{4D_1\sigma^L}} e^{\sqrt{\eta_1} y} dy \right) e^{-\sqrt{\eta_1}x}, \qquad (38)
\end{aligned}$$

which is easily seen to still be monotonically decreasing for all $x \geq 0$ since $\frac{d}{dx}\left(\frac{1}{\sqrt{\eta_1}} + x\right) e^{-\sqrt{\eta_1}x} = -\sqrt{\eta_1} x e^{-\sqrt{\eta_1}x} \leq 0$.

In Fig. S1, we confirm that a high $\beta_2^N$ is in fact needed to create the non-monotonic gradients we observe in $\overline{N_Q^i}(x)$ inside the exclosure, showing at least empirically that this same result holds in the case of the circular exclosure. We have thus shown that the non-monotonic gradient created by questing tick movement occurs because questing ticks closer to the edge of the exclosure are more likely to journey outside of the exclosure and be picked up by deer. This result supports the "tick sharing" mechanism hypothesized in [6, 8] to explain why tick densities are reduced at the inside edge of an exclosure. In other words, as we increase $\beta_2^N$, we reduce $\overline{N_Q^i}(x)$ everywhere inside the exclosure, but this reduction is more dramatic for small positive $x$ than for large positive $x$; and when $\beta_2^N$ is sufficiently high, we can then create a non-monotonic gradient of $\overline{N_Q^i}(x)$ inside the exclosure. We make this argument more rigorous in the next result. While we are not able to draw as nice of analytic results when $\beta_2^N > 0$, we are able to draw this general conclusion:

**Theorem 3.** *Assume $\beta_1^A = 0$, $\alpha = 0$, and $D^L = 0$ but $D^N > 0$. Then the gradient of $\overline{N_Q^i}(x)$ for $x \geq 0$ will be non-monotonic if and only if $\beta_2^N$ is sufficiently large. And no matter the value of $\beta_2^N$, the gradient $\overline{N_Q^i}(x)$ as $x \to \infty$ converges to the function given by (34). Consequently, $\overline{N_Q^i}(x)$ for large $x$ converges to an exponential decay at rate $\min\{\sqrt{\eta_1}, \sqrt{\eta_2}\}$, i.e.*

$$\lim_{x\to\infty} \frac{d}{dx} \log \overline{N_Q^i}(x) = -\min\left\{\sqrt{\frac{\sigma^L}{D_1}}, \sqrt{\frac{\beta_1^N K_1 + d^N}{D^N}}\right\} \tag{39}$$

*Proof.* With $\beta_2^N > 0$, (35) no longer applies. Instead, we get the more general expression

$$\overline{N_Q^i}(x) = \int_{-\infty}^{\infty} \int_0^{\infty} \frac{m^L \sigma^L \overline{L_{F_1}^e}(y)}{\sqrt{4\pi D^N \tau}} e^{-\frac{(x-y)^2}{4D^N\tau}} p(x, y, \tau, \beta_1^N K_1 + d^N, \beta_2^N K_2, D^N) d\tau dy \quad \forall x \in \mathbb{R}, \tag{40}$$

where $p(x, y, \tau, \delta_1, \delta_2, D)$ is the probability that a Brownian bridge $W_{y,t=0}^{x,t=\tau}(t)$ with diffusion coefficient $D$ starting at $y$ and ending at $x$ after time $\tau$ "survives" until $t = \tau$, given that it "fails" at rate of $\delta_1$ whenever $W_{y,t=0}^{x,t=\tau}(t) \geq 0$ and at a rate of $\delta_1 + \delta_2$ whenever $W_{y,t=0}^{x,t=\tau}(t) < 0$. This quantity can be expressed as

$$p(x, y, \tau, \delta_1, \delta_2, D) = e^{-\delta_1 \tau} \int_0^{\tau} e^{-\delta_2 \zeta} \, \mathbb{P}\left[\int_0^{\tau} \mathbb{1}\left\{W_{y,t=0}^{x,t=\tau}(t) < 0\right\} dt \in (\zeta, \zeta + d\zeta)\right], \tag{41}$$

where $\mathbb{1}\{X\}$ is an indicator function equal to 1 when $X$ is true and 0 otherwise. When $\beta_2^N = 0$, we see that $p(x, y, \tau, \beta_1^N K_1 + d^N, \beta_2^N K_2, D) = e^{-\delta_1 \tau}$, and so (40) will reduce to (35) (resulting in a monotonic gradient for $\overline{N_Q^i}(x)$ for $x \geq 0$ by Theorem 2). But when $\beta_2^N > 0$, there is no nice analytic expression to our knowledge for $\mathbb{P}\left[\int_0^{\tau} \mathbb{1}\left\{W_{x,t=0}^{y,t=\tau}(t) < 0\right\} dt \in (\zeta, \zeta + d\zeta)\right]$ (the distribution for the amount of time a Brownian bridge with general start and end points spends below 0).

In the limit as $\delta_2 \to \infty$, however, we are able to derive an expression for $p(x, y, \tau, \delta_1, \delta_2, D)$, using the fact that $\mathbb{P}\left[\min_{0 \leq t \leq \tau} W_{y,t=0}^{x,t=\tau} \geq 0\right] = 1 - e^{-\frac{xy}{D\tau}}$ when $x, y \geq 0$ [9]. When $\delta_2$ is infinitely large, the Brownian bridge is guaranteed to "fail" the instant it passes below 0, and so the expression inside the outer integral of (41) is only non-zero when $\zeta = 0$. Consequently, the outer integral of (41) will evaluate to the probability that the Brownian bridge never goes below 0 for $t \leq \tau$, which equals $1 - e^{-\frac{xy}{D\tau}}$ when $x, y \geq 0$ and equals 0 when $x < 0$ or $y < 0$. Thus,

$$\lim_{\delta_2 \to \infty} p(x, y, \tau, \delta_1, \delta_2, D) = \begin{cases} e^{-\delta_1 \tau}\left(1 - e^{-\frac{xy}{D\tau}}\right) & \text{if } x, y \geq 0, \\ 0 & \text{otherwise.} \end{cases} \tag{42}$$

In the limit as $\beta_2^N \to \infty$, we can plug this result into (40) to derive for $x \geq 0$

$$\begin{aligned}
\overline{N_Q^i}(x) &= \int_0^\infty \int_0^\infty \frac{m^L \sigma^L \overline{L_{F_1}^e}(y)}{\sqrt{4\pi D^N \tau}} e^{-\frac{(x-y)^2}{4D^N\tau}} e^{-(\beta_1^N K_1 + d^N)\tau} \left(1 - e^{-\frac{xy}{D^N\tau}}\right) d\tau dy \\
&= \int_0^\infty \int_0^\infty \frac{m^L \sigma^L \overline{L_{F_1}^e}(y)}{\sqrt{4\pi D^N \tau}} e^{-\frac{(x-y)^2}{4D^N\tau}} e^{-(\beta_1^N K_1 + d^N)\tau} d\tau dy \\
&\quad - \int_0^\infty \int_0^\infty \frac{m^L \sigma^L \overline{L_{F_1}^e}(y)}{\sqrt{4\pi D^N \tau}} e^{-\frac{(x+y)^2}{4D^N\tau}} e^{-(\beta_1^N K_1 + d^N)\tau} d\tau dy \\
&= \frac{m^L \sigma^L}{\sqrt{4D^N(\beta_1^N K_1 + d^N)}} \left[\int_0^x \overline{L_{F_1}^e}(0^+) e^{-\sqrt{\eta_1} y} e^{-\sqrt{\eta_2}(x-y)} dy + \int_x^\infty \overline{L_{F_1}^e}(0^+) e^{-\sqrt{\eta_1} y} e^{-\sqrt{\eta_2}(y-x)} dy \right. \\
&\quad \left. - \int_0^\infty \overline{L_{F_1}^e}(0^+) e^{-\sqrt{\eta_1} y} e^{-\sqrt{\eta_2}(x+y)} dy \right] \\
&= \frac{m^L \sigma^L}{\sqrt{4D^N(\beta_1^N K_1 + d^N)}} \left[\frac{\overline{L_{F_1}^e}(0^+)}{\sqrt{\eta_2} - \sqrt{\eta_1}} \left(e^{-\sqrt{\eta_1}x} - e^{-\sqrt{\eta_2}x}\right) + \frac{\overline{L_{F_1}^e}(0^+)}{\sqrt{\eta_1} + \sqrt{\eta_2}} e^{-\sqrt{\eta_1}x} \right. \\
&\quad \left. - \frac{\overline{L_{F_1}^e}(0^+)}{\sqrt{\eta_1} + \sqrt{\eta_2}} e^{-\sqrt{\eta_2}x} \right] \\
&= \frac{m^L \sigma^L}{\sqrt{4D^N(\beta_1^N K_1 + d^N)}} \left(\psi_1 e^{-\sqrt{\eta_1}x} - \psi_1 e^{-\sqrt{\eta_2}x}\right),
\end{aligned} \tag{43}$$

restricting the outer integral in the first line to $y \geq 0$ since $p(x, y, \tau, \delta_1, \delta_2, D) = 0$ for $y < 0$, applying Lemma 1 and (32), and with $\psi_1$ defined in (34) (note the close similarity of (43) to (34)). Now if we have $\eta_1 < \eta_2$, then $\psi_1 > 0$, so applying Lemma 2 to (43), we get that $\overline{N_Q^i}(x)$ is non-monotonic for $x \geq 0$, with an initial increase followed by a decline to 0. Similarly, if $\eta_1 > \eta_2$, we can apply Lemma 2 with $-\psi_1 > 0$ to get that $\overline{N_Q^i}(x)$ is still non-monotonic for $x \geq 0$. (As before, we ignore the case where $\eta_1 = \eta_2$, as it is impossible for the parameters satisfy this perfectly in practice, though we address it in another remark after this proof). Again, this result of a non-monotonic gradient comes from assuming a very large $\beta_2^N$ approaching $\infty$, while assuming $\beta_2^N = 0$ leads to a monotonically decreasing gradient of $\overline{N_Q^i}(x)$ for $x \geq 0$ by Theorem 2. Thus, by continuity, the gradient must be monotonically decreasing for any sufficiently small $\beta_2^N$ by continuity, and the gradient must be non-monotonic for any sufficiently large $\beta_2^N$.

No matter the value of $\beta_2^N$ or the value of $y$, it must be true that

$$\lim_{x \to \infty} p(x, y, \tau, \delta_1, \delta_2, D) = e^{-\delta_1 \tau}. \tag{44}$$

This comes from the fact that, as $x$ gets very large, a Brownian bridge conditioned to end at $x$ after a fixed amount of time must spend very little time below 0; and as $x$ approaches infinity with everything else fixed, the time spent below 0 will go to 0. As such, the expression inside the outer integral of (41) will only be non-zero when $\zeta = 0$, providing us with (44). Thus, as $x \to \infty$, (40) will reduce to (35), from which the rest of the proof of Theorem 2 shows that the the gradient of $\overline{N_Q^i}(x)$ will approach that given by (34). And applying Lemma 2 to (34) shows that $\overline{N_Q^i}(x)$ will have an asymptotic exponential decay rate of $\min\{\sqrt{\eta_1}, \sqrt{\eta_2}\}$ as $x \to \infty$.

□

**Remark.** Empirically, it will generally be the case in practice that $\frac{\sigma^L}{D_1} < \frac{\beta_1^N K_1 + d^N}{D^N}$, since rodents generally have significantly more movement than ticks, leading to $D_1 \gg D_N$. This is exacerbated by the fact that an animal's diffusion coefficient scales quadratically with its average daily movement rate. Thus, the asymptotic rate of exponential decay for $\overline{N_Q^i}(x)$ as $x \to \infty$ will most likely $\sqrt{\frac{\sigma^L}{D_1}}$ in realistic scenarios, the same as it is for the case with no questing tick movement in Theorem 1.

**Remark.** In the case of $\eta_1 = \eta_2$, using L'Hôpital's rule in the limit as $\zeta = \eta_2 - \eta_1 \to 0$ on (43) for $x \geq 0$ yields

$$\begin{aligned} \overline{N_Q^i}(x) &= \frac{m^L \sigma^L}{\sqrt{4D^N(\beta_1^N K_1 + d^N)}} \lim_{\zeta \to 0} \frac{2\sqrt{\eta_1 + \zeta}\,\overline{L_{F_1}^e}(0^+)\left(e^{-\sqrt{\eta_1}x} - e^{-\sqrt{\eta_1+\zeta}x}\right)}{\zeta} \\ &= \frac{m^L \sigma^L}{\sqrt{4D^N(\beta_1^N K_1 + d^N)}} \overline{L_{F_1}^e}(0^+) x e^{-\sqrt{\eta_1}x}, \end{aligned} \tag{45}$$

which is non-monotonic over $x \geq 0$.

#### 5 Visual evidence that a non-monotonic $\overline{N_Q^i}$ gradient results if and only if $\beta_2^N$ is sufficiently large

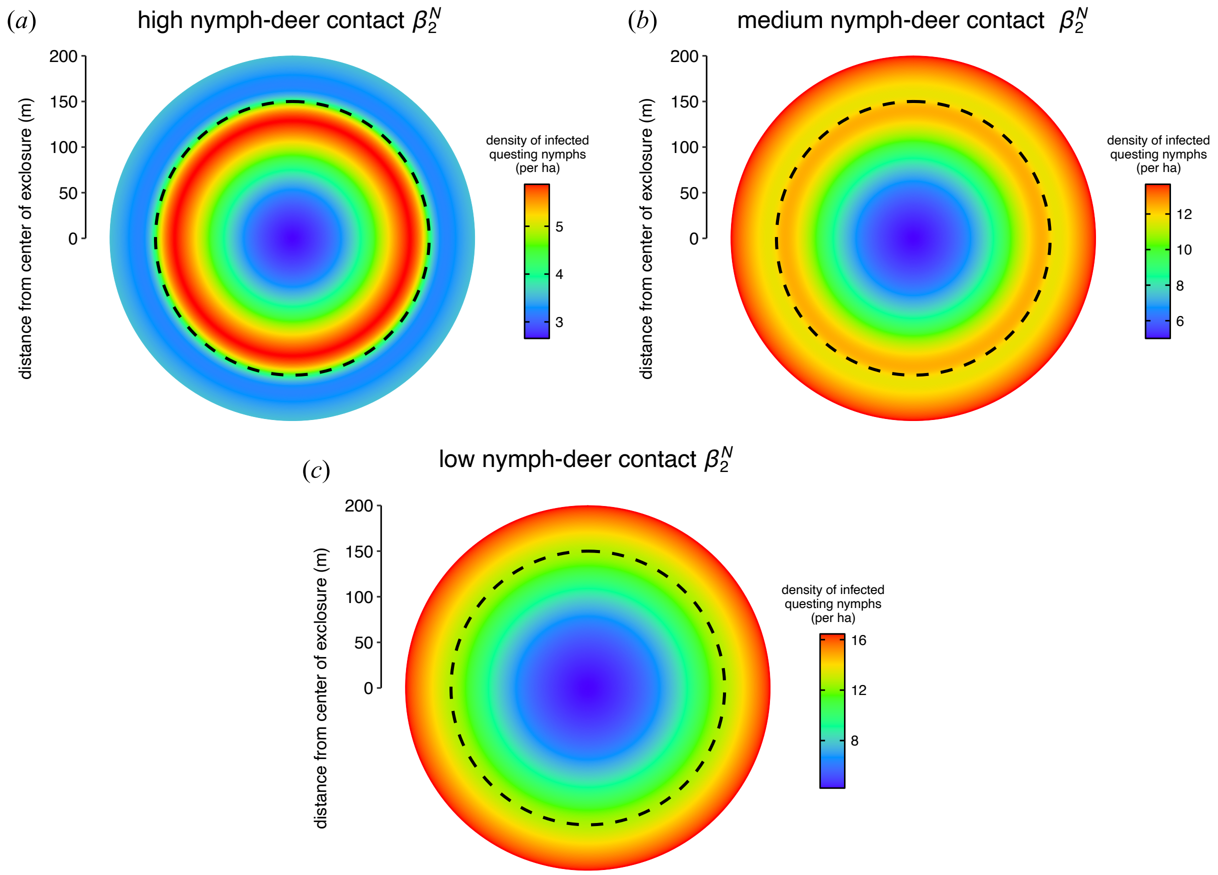

Figure S1: Infected questing nymph density at equilibrium across space with questing tick movement and a large exclosure (radius of 150 m, 7.1 ha), simulated with the same parameters as Fig. 5b in the main text but with varying levels of $\beta_2^N$, the nymph-deer contact rate. In (a), $\beta_2^N$ is the same as it is for Fig. 5b in the main text; in (b), $\beta_2^N$ is reduced to 25% of this value; and in (c), $\beta_2^N$ is reduced to 10% of this value. Note that the color scale bars in $(a)$–$(c)$ are all different.

This shows that Theorem 3 holds in simulation even with a large circular exclosure and a small degree of infection-based mortality: (a) has a sufficiently high $\beta_2^N$, resulting in a non-monotonic gradient inside the exclosure, while $\beta_2^N$ is reduced in (b) and (c) to the point where the gradient becomes monotonically decreasing. Reducing $\beta_2^N$, however, comes at the loss of the dilution effect: in (b), there is still a small dilution effect creating slightly higher infected questing nymph density at the inside edge of the exclosure than the outside edge, but in (c) the dilution effect is completely lost as the density is lower everywhere in side the exclosure than anywhere outside. This happens because nymphs contacting with deer is essential to producing the dilution effect in the first place, as deer are dilution hosts [10], and reducing $\beta_2^N$ reduces nymphs' contact with deer by definition.
